# Matchathon: A guide to student-faculty connections in PhD programs

**DOI:** 10.1101/2020.11.06.371526

**Authors:** Haley Amemiya, Zena Lapp, Cathy Smith, Margaret Durdan, Michelle DiMondo, Beth Bodiya, Scott Barolo

**Affiliations:** Cellular and Molecular Biology Program, University of Michigan Medical School, Ann Arbor, MI, USA; Department of Bioinformatics and Computational Biology, University of Michigan Medical School, Ann Arbor, MI, USA; Program in Biomedical Sciences, University of Michigan, Ann Arbor, MI, USA; Office of Graduate & Postdoctoral Studies, University of Michigan, Ann Arbor, MI, USA; York College of Pennsylvania, York, PA, USA; Department of Cell and Developmental Biology, University of Michigan Medical School, Ann Arbor, MI, USA

## Abstract

Relevant and impactful mentors are essential to a graduate student’s career. Finding mentors can be challenging in umbrella programs with hundreds of faculty members. To foster connections between potential mentors and students with similar research interests, we created a Matchathon event, which has successfully enabled students to find mentors. We developed an easy-to-use R Shiny app (https://github.com/UM-OGPS/matchathon/) to facilitate matching and organizing the event that can be used at any institution. It is our hope that this resource will improve the environment and retention rates for students in the academy.

The open source app is publicly available on the web (app: https://UM-OGPS.shinyapps.io/matchathon/; source code: https://github.com/UM-OGPS/matchathon/).

## Introduction

Graduate level attrition rates, especially for doctoral candidates, are one of academia’s highest; estimated to be from 40-50% [1,2]. The prominent theory for such high attrition is the failure of social, professional, and academic integration of students into their department [1],[3]. When surveyed, students cite a disconnect between their research mentor and themselves regarding research goals and expectations [1],[4],[5] and inadequate reassurance on their personal dissertation and research progress from thesis committee members. Furthermore, successful advisor-student relationships influence downstream success such as productivity, mental health, and achieving career goals [6]. However, it can be difficult for incoming students to identify prospective research mentors, particularly in large “umbrella” programs where students can have a mentor in any of several different departments. For instance, students in the University of Michigan’s (UM) Program in Biomedical Sciences (PIBS) have access to 500+ faculty mentors across 14 PhD programs, and can perform rotations in three to six labs. Faculty with similar research interests may not be on the students’ radar due to the overwhelming number of faculty researchers on campus, and because faculty websites may contain outdated research interests. Furthermore, each year only a fraction of faculty have funding and are actively looking for students.

One method that some umbrella programs have implemented to expose students to potential faculty mentors is faculty talks; however, students are often disengaged, few faculty speak at these events, and sometimes not all faculty speakers have funding for new students. To facilitate potential student-mentor connections, we created a matching algorithm to pair students and faculty based on common research interests, and established a structured “Matchathon” event where students meet multiple faculty members individually to discuss shared research interests (Figure 1). We hosted Matchathon in person for three years (2017-2019); however, due to the COVID-19 pandemic, we moved to a virtual platform in 2020. The Matchathon event helps incoming doctoral candidates identify potential advisors, mentors, and committee members who can provide encouragement and guidance throughout the dissertation process.

**Figure 1:**
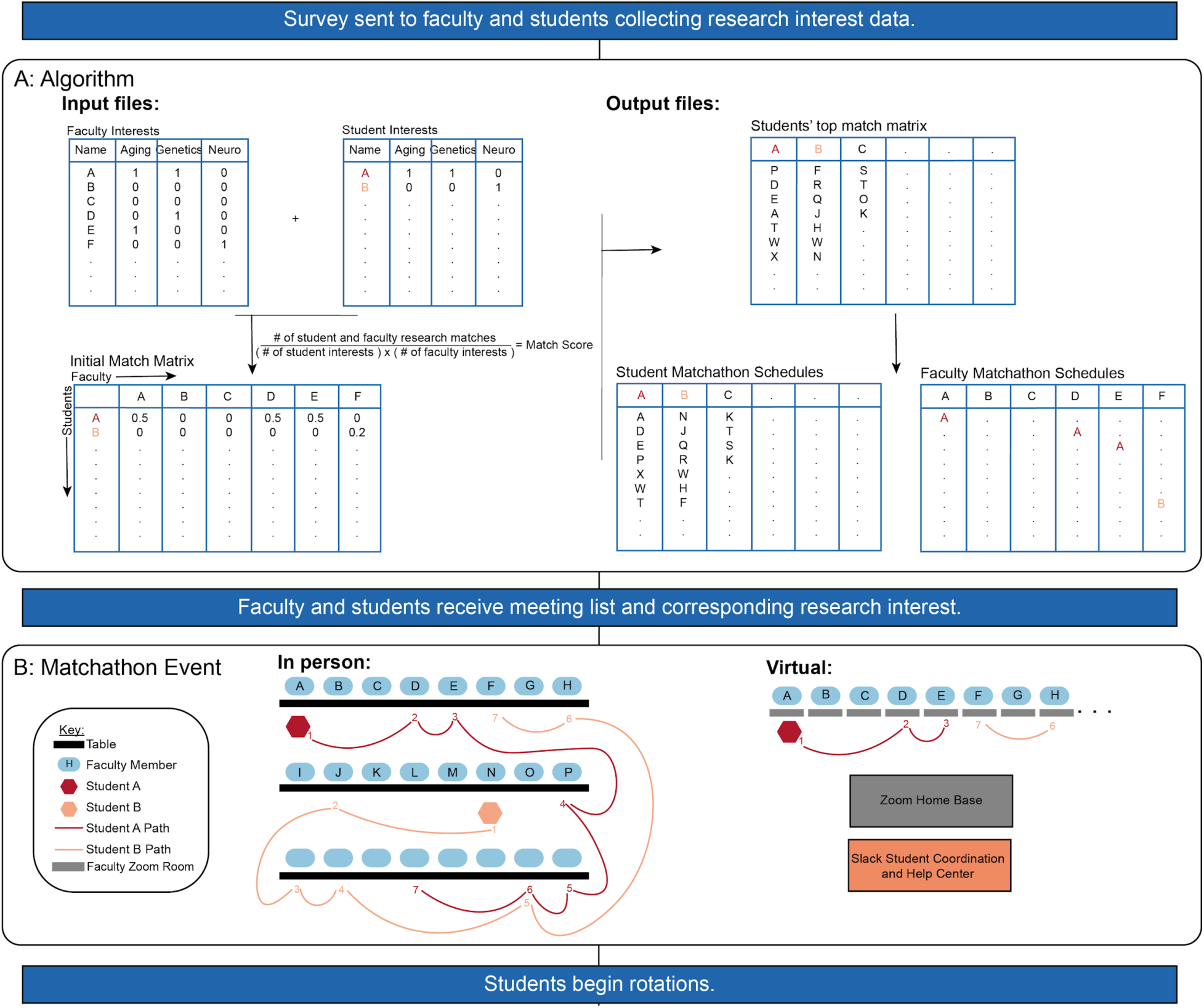
Matchathon algorithm and event timeline. (A) Algorithm overview. Input files must have the same column names to create a match score based on research interest overlap. These scores are used to rank student-faculty matches, and create a schedule for the event. (B) Example path of two students (A (red) and B (tan)) during Matchathon for in person vs virtual platforms. For the in-person set up, faculty remain at their post while students rotate every seven minutes. In the virtual platform, each faculty member has their own Zoom room. A link to a Zoom home base was provided to both faculty and students where a team of organizers assisted with technical difficulties. Lastly, Slack was utilized as another method to communicate with students during Matchathon.

## General overview of Matchathon

To help students find faculty members with mutual research interests, we use an algorithm to match first-year PhD students and faculty who are in search of trainees based on mutual research interests (Figure 1A). The algorithm and Matchathon schedule-builder are packaged in an easy-to-use open source R Shiny app [7], and the corresponding Matchathon package can be downloaded from GitHub (https://github.com/UM-OGPS/matchathon/) or used on the web (https://UM-OGPS.shinyapps.io/matchathon/).

During Matchathon, students meet individually with 12 different faculty members (Figure 1B). From 2017 to 2019 Matchathon was an in-person event; however, in 2020 we organized the first virtual Matchathon using the Zoom video conferencing platform [8]. While both events are described here, the administration, faculty, and students preferred the virtual event, largely due to a less noisy environment. Detailed information about the algorithm, communicating the event to faculty and students, and running both the in person and virtual events, can be found in the supplemental material.

## Results

We have conducted Matchathon for the past four years (2017-2020). Students from 2017 to 2019 have already chosen labs. Furthermore, in 2019 and 2020 we asked students to list the names of any faculty members they had already met with to maximize new student-faculty connections at the event. We found that both students and faculty benefitted from Matchathon.

### Matchathon helps students find rotation and thesis labs

To determine whether Matchathon helps students identify potential research mentors, we surveyed students from the 2017, 2018, and 2019 incoming cohorts about their Matchathon experience. For the years 2017-2019, 34 students found a rotation mentor at Matchathon (Figure 2A). Note that the increase in students who found a rotation through Matchathon in 2019 may be because we ensured that students did not meet with faculty members they were already in touch with; however, we would need more longitudinal data to confirm this finding. Furthermore, Matchathon helped 14 students find a PhD mentor (Figure 2B). While we realize that these numbers might sound small, finding the correct mentor is extremely important for success in graduate school, and our results indicate that 14 graduate students joined a lab that they might not have found if Matchathon did not happen. One graduate student explained:

**Figure 2:**
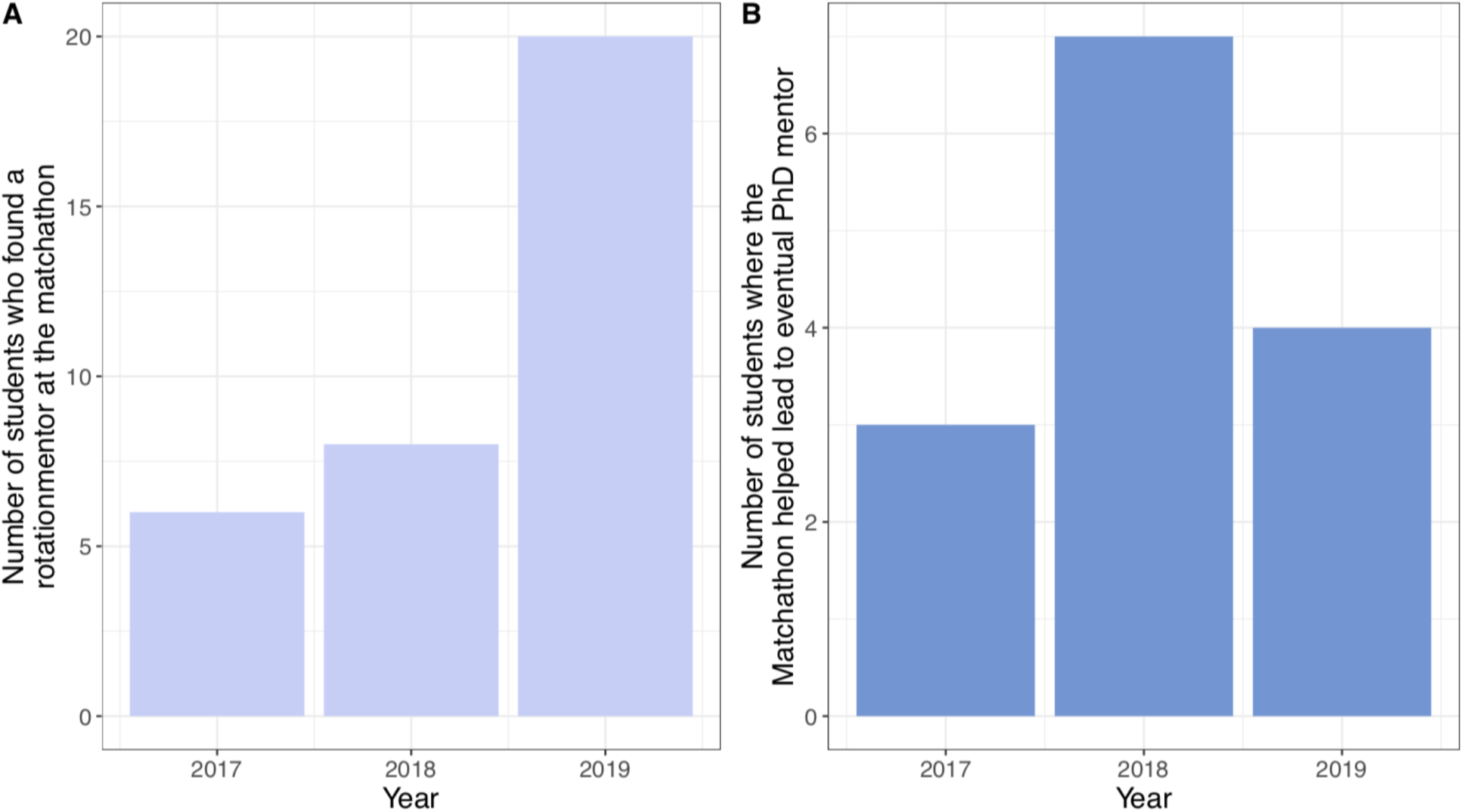
Matchathon efficacy. Based on the survey results: (A) number of students who found a rotation mentor at Matchathon, and (B) number of students where Matchathon helped lead them to their eventual PhD mentor.

> The matchathon led me to rotate in a laboratory I hadn’t heard of before. Over the course of the half rotation I was able to narrow in my research interests and find an incredible mentor and some great new friends. I fully intend on joining this laboratory now and it would have never happened without the matchathon.

More quotes from students about the event can be found in the supplementary material.

### Faculty find Matchathon useful for setting up new rotations

While student participation in Matchathon is required (cohort size each year: 72, 81, 97, and 87, respectively), faculty participation is voluntary. There was a stark increase in faculty participation after the first year (number of participants each year: 57, 93, 106, and 108, respectively). Furthermore, across all years, 28/87 (32%) of the faculty who filled out the survey claimed that Matchathon helped them set up new rotations in their lab. Although the focus of the event is on students benefiting from Matchathon, this suggests that faculty members may also find the event helpful for identifying potential student trainees. Furthermore, feedback from faculty, particularly about the virtual event, is quite positive (see supplementary material).

## Discussion

Here we present Matchathon and show that it is a useful way for students and faculty with similar research interests to connect. Students do not have to prepare for the event, and it puts the pressure on faculty to explain their research and lab to students in an engaging way. Students meet with faculty members who can later become their rotation and thesis mentors, and the number of faculty participating has increased since the first year. Furthermore, we have received positive feedback from both students and faculty about the event, and increasingly positive feedback about the virtual format over the in-person event. Thus, events such as these may improve graduate student retention rates and overall graduate student satisfaction.

## Methods

### Matchathon algorithm & logistics

#### Matching students with faculty

Prior to the event we collect a list of faculty with funding and openings for new graduate students. This ensures that students are only meeting faculty members recruiting new trainees (i.e. taking rotation students for the upcoming year). We then provide a survey to both students and faculty that includes a list of research topics within the program (example surveys: https://drive.google.com/drive/folders/1h5t1tlCVsan0vAjtq1EgioJmd_fghYT6?usp=sharing). Faculty members select interests that pertain to their lab, and before arriving at UM for graduate school, the students select their research interests. Additionally, to promote new student-faculty connections, we ask students to provide a list of faculty members they have already met with, and ensure they are not matched for the event. We then use the data from the surveys as inputs to our algorithm.

#### Matching algorithm & creating Matchathon schedule

After the research interest collection is finalized, we feed the files into the algorithm (Figure 1A). The first step counts the number of mutual research interests for each student-faculty pair, as well as the total number of interests of the pair. These components (the number of overlaps, total faculty interests, and total student interests) are used to calculate a match score for each faculty and student pair using the following equation:

\# of student and faculty research matches / (# of student interests * # of faculty interests)

The overall match score is smaller for those who selected a high number of interests, thus prioritizing matches where research interests are more focused. We then rank the scores for each student to generate a list of the top matched faculty in descending order. This ranked file is what we use to generate the schedule with a user-defined number of time slots (12 in our case). We exclude faculty members that students have already met, and require that faculty must have at least half of their schedule full. Additionally, if certain faculty members must leave early or arrive late, this information can be included as input to the algorithm, and faculty will be assigned as unavailable for those time slots.

The algorithm and Matchathon schedule-builder are packaged in an easy-to-use open source R Shiny app, and the corresponding Matchathon package can be downloaded from GitHub (https://github.com/UM-OGPS/matchathon/) or used on the web (https://UM-OGPS.shinyapps.io/matchathon/).

#### Matchathon

Matchathon took place in person in 2017, 2018, and 2019, and virtually over Zoom in 2020. The coordinators give a brief introduction to the event, and then the students meet with 12 different faculty members. For the in person event, for 5 minutes with each faculty, and 2 minutes of transition time between meetings, while the virtual event allowed for 7 minute meetings, with 3 minutes of transition time. We also include a 10 minute break after half of the meetings are over. See the supplementary material for more details about running the in-person and virtual events.

#### Post-matchathon survey

Using Google Forms or Qualtrics, we surveyed the students and faculty who participated in Matchathon after each annual event. The brief surveys vary slightly from year to year, but generally, faculty and students respond about demographic information and report their level of satisfaction with Matchathon. See supplementary material for more information about the survey content. The results from these surveys are compiled to evaluate the efficacy of Matchathon.

#### R version & packages

R version 3.6.2 [9] was used to create the R Shiny app. The following packages were used in the R Shiny app and/or to generate figures: shiny version 1.4.0 [7], tidyverse version 1.3.0 [10], randomNames version 1.4-0.0 [11,12], readxl version 1.3.1 [11], cowplot version 1.0.0 [7,13].

#### Communicating with students about the event

Students in PIBS accept their offer of admission to the program in mid-April and matriculate into the program either in summer (July) or the fall (late-August). Matchathon is advertised to incoming PIBS students as early as their first email from the PIBS Director and Program Manager. This email provides them an introduction into the program and lists all of the upcoming events for first-year students. Matchathon is also mentioned during several onboarding emails to the incoming PIBS cohort. Specifically, in mid-June students are emailed a list of resources for finding potential rotation matches. In this email, Matchathon is introduced to the PIBS cohort.

Incoming students receive a follow-up message in early July asking them to fill out a Google Form about their research interests. This allows PIBS to capture any changes in research interests from when they applied to the program the prior fall. In this form, we allow students to opt-out of meeting with faculty whom they already know and opt-out of meeting with faculty whom they have already decided to do a rotation with. In August before Matchathon, students are emailed detailed information about the event. The list of faculty matches is provided, which allows the student to (optionally) research the faculty matches prior to the event.

#### Communicating with faculty about the event

Faculty research mentors affiliated with PIBS are contacted in May about their interest in taking PIBS students for rotations and their interest in attending Matchathon. Faculty are not required to attend Matchathon and may opt-out of participating; however if a faculty member is available and interested in attending, the PIBS Director asks them to update their research interests via a Google Form. We also remind them to add the date of the event to their calendars and we let them know that they will receive more information (i.e. the location and their student matches) in mid-August. In mid-August faculty are emailed a reminder of the event and given a link to a Google sheet that lists their student matches. We send this email to faculty before we email the incoming PIBS cohort because schedules for faculty change quickly and it is normal for a handful of faculty members (∼5 to 15) to drop out of the event last minute. When this happens the matching algorithm is updated to remove those faculty members.

#### Running the in-person event

Due to the size of this event (200+ faculty and students) finding the right venue early on is important. The event space needs to be a large room with movable tables and chairs. Once the space is reserved it is necessary to order any additional supplies or materials to host the event. At UM these supplies include items for hosting (water, tea, and coffee), event equipment (tables, table cloths, and chairs), and office supplies (table tents with faculty names, name tags for students and faculty, place card holders identifying the alphabet, scratch paper, pens, and hard copies of the student meeting agenda). We also create a staff agenda to identify staff tasks and placement to make sure the event runs smoothly. We have staff who escort our students and faculty to the event space. We have staff who help students find their faculty matches. There also is a staff member who times each of the seven minute meetings and notifies students and faculty when it is time to end conversations and move on to the next match.

#### Running the virtual event

In a similar fashion to the in-person Matchathon, the virtual format of the event involves reaching out to students and faculty prior to the event. This includes informing students about the event and sending information to faculty members about participation, as well as the research interest survey. When the surveys are submitted, the algorithm is run, schedules produced, and each schedule is sent out to individuals via mail merge. The same logistics regarding the generation of schedules apply to the virtual format. About 1-2 weeks prior to the virtual event an email is sent to the faculty members with instructions on how to set up a Zoom link to a personal Zoom room. The instructions include how to create their room, and how to take off password requirements and the waiting room option. This is to ensure students entering a new meeting do not have to wait to be “let in” to the meeting, which also helps move meetings along. Individual faculty members remain in their personal Zoom room throughout the event.. The Zoom room link and/or meeting ID that is provided by faculty members is included on the student schedule, and students have the responsibility to enter and leave each meeting. Facilitators of the event coordinate a Zoom meeting just before the meetings begin to kick-off the event and remind both students and faculty about the logistics. The facilitators remain on that Zoom call so that students or faculty can return to the room if problems arise.

In addition to helping individuals in the Zoom Home Base, a Slack channel is used as a helpline to address and respond to technical, Zoom, or general issues. Students are able to reach out to the facilitators in live-time. Slack is also used to send time notifications to students so that they do not have to keep an extensive watch on the clock, and can wait for a Slack message to notify them that it is time to move on to the next meeting. Slack is used for other announcements as well, including reminders about how many meetings are left, and notification of a short break between meetings. We provided a 10 minute break halfway through the meetings, similar to the in-person event. Following the event, surveys are sent to students and faculty members to collect feedback about the event and how it can be improved.

#### Post-matchathon surveys

Both faculty and student surveys begin with demographic questions (main department affiliation, year they joined the university, and, for the student survey only, a validation question to affirm they participated in Matchathon during the current year). Once we validate their participation in Matchathon, we then ask students about their satisfaction with the event. For the student survey, participants report how many rotations they had set up before the event (range 0-5). They then indicate which potential benefits they experienced from their Matchathon participation (ie ‘Matchathon was valuable for networking and connecting with faculty’, ‘Matchathon helped connect me with my PhD mentor’). Finally, there are several free response options to share any other feedback from the event (‘What would you change about Matchathon?’). The satisfaction section of the faculty survey is similar, however since faculty can participate across multiple years, they indicate any barriers to their participation in the event (childcare, time conflicts, lack of understanding of the purpose of the event, and for 2020, technological issues). Faculty then report on both their positive and potential negative experiences with the event (ie ‘Matchathon helped to set up new rotations in my lab’. ‘Matchathon matched me with students who didn’t share my research interests’). Faculty indicate if the event fostered longer meetings with students later (yes/no), and for 2020 only, faculty members who participated across multiple years report their preference for the virtual or in person event (preferred virtual/preferred in person/no preference). The faculty survey then ends with free response boxes to share any other feedback or suggestions for future events (‘What aspects of Matchathon 2020 could be improved in the future?’).

#### Positive feedback on the event and the virtual format

The transition to a virtual event in the fall of 2020 was supported by faculty, students, and facilitators, and the faculty overwhelmingly recommended us to continue using the virtual platform in subsequent years. Feedback from students and faculty from the in-person event included:

> “*Matchathon is great, and is very useful for connecting student and faculty*”
>
> “*The format was fun and I liked that I connected with faculty I never would’ve met outside of this event*.”

A reoccurring suggestion for making the event better included comments that felt the space used for the event was too crowded, including “*Ultimately, an amazing event-I speak very highly of it-just needs to be in a bigger space*” and “*I would keep everything the same, but it needs to be in a quieter setting*”. From the Fall 2020 survey, we received a great majority of participants that preferred the virtual format. Faculty expressed their appreciation and support in continuing online:

> “*The matchathon is so much better this year! Zoom is definitely the way to go*” “*I think this was better than the in-person version*”
>
> “*The reduction of noise makes this so much better. Overall calmer and easier to talk. I would vote to keep this system for the future*.”
>
> “*It was even more productive than the in-person meetings of the past!*”
>
> “*never go back to in person match a thon*”
>
> “*Holding the meeting in a single room with 100s of participants make it far too loud and crowded; the virtual event solves this and, strangely, makes it more personal*”

Overall, the overwhelming majority of participants have given positive feedback for both the in-person and virtual events.

> “*Really worked well to help me introduce myself to some very meaningful connections/students*.”
>
> “*The range of research interests was phenomenal*”
>
> “*The matching system was really good, I was matched to quite a few people who I hadn’t heard of but ended up finding very interesting*”
>
> “*The matching program or algorithm used was extremely effective. I am currently rotating with a faculty I met through the matchathon event. It also made me realize how diverse my interests are as none of the faculties that I was matched with was in my department and therefore the chances of coming across their research were very slim if it wasn’t for the matchathon*.”
>
> “*The matchathon led me to rotate in a laboratory I hadn’t heard of before. Over the course of the half rotation I was able to narrow in my research interests and find an incredible mentor and some great new friends. I fully intend on joining this laboratory now and it would have never happened without the matchathon*”
>
> “*It was a great experience and idea. I am grateful I could participate in it*.”

## Author Contributions

Conceptualization: H.A., Z.L., M.D., and S.B.; Methodology: H.A., Z.L., M.D., and S.B.; Software: H.A., Z.L., C.S., M.M.D.; Formal analysis: H.A., Z.L.; Investigation: H.A., Z.L.; Visualization: H.A., Z.L.; Supervision: M.D. and S.B.; Project Administration: M.D. and S.B.; Writing - Original Draft: H.A., Z.L., M.D., S.B., ; Writing – Review & Editing: H.A., Z.L., C.S., M.M.D., M.D., B.B., S.B.

